# Prebiotic aqueous reactions catalyzed by native nickel without hydrogen

**DOI:** 10.64898/2026.01.05.697669

**Authors:** Carolina Garcia Garcia, Max Brabender, William F. Martin

**Author notes:** Author for correspondence: Carolina Garcia Garcia, Institute for Molecular Evolution, Düsseldorf University, Universitätsstrasse 1, 40225 Düsseldorf, Germany.

## Abstract

Compared to iron, nickel is comparatively rare as a transition metal in enzymes. But it is essential in several enzymes of carbon and energy metabolism in acetogens (bacteria) and methanogens (archaea), which use the acetyl-CoA pathway of H_2_-dependent CO_2_ fixation. Nickel containing enzymes of acetogens and methanogens include FeNi hydrogenase, carbon monoxide dehydrogenase, acetyl-CoA synthase and, in methanogens, methyl-CoM reductase in the last step of methane synthesis. Several lines of evidence implicate the acetyl-CoA pathway as the most ancient pathway of CO_2_ fixation, most notably recent findings that the overall reaction of the enzymatic pathway from H_2_ (*E*_0_′ = –414 mV) and CO_2_ to pyruvate can be replaced by Ni^0^ alone in water as the lone catalyst. Here we studied the ability of Ni^0^ to serve as catalyst and reductant for nonenzymatic redox reactions that require only a mild reductant, as the midpoint potential of Ni^0^ oxidation to Ni^2+^ is *E*_0_′ = –260 mV. We show that Ni^0^ in water can convert 2-oxo acids to 2-hydroxy acids and, in the presence of NH_3_, to amino acids at 25-100°C without addition of H_2_, and that it will function as catalyst and reductant for the fumarate reductase reaction. The findings expand the repertoire of ancient metabolic reactions that Ni^0^ can catalyze without proteins, cofactors, or sulfur, shedding light on the broad catalytic activity and substrate specificity of Ni^0^ at metabolic origin.

## INTRODUCTION

Metabolism is a chemical reaction. It emerged from reactions catalyzed by environments present on the early Earth and gave rise to the metabolic reaction networks of living cells (1–4). Modern microbes possess enzymes and cofactors that accelerate the reactions of metabolism so that all reactions take place at roughly the same rate (5). At the very onset of metabolism, there were no enzymes or cofactors, only inorganic catalysts (6). Transition metals play a crucial role in theories for metabolic origin because they are notoriously good at accelerating particularly difficult chemical reactions such as N_2_ reduction (7). Coordinated in proteins and cofactors, Fe, Ni, Co, and Mo (sometimes replaced by W (8)), are essential to catalysis in anaerobic autotrophs and are abundant in enzymes of the acetyl-CoA pathway (9–14). Among pathways of CO_2_ fixation, the acetyl-CoA pathway is the most ancient, the only one that occurs in bacteria and archaea (15–16) and the only one that serves both carbon and energy metabolism (13,18). It is the starting point of metabolism in acetogens (bacteria) and methanogens (archaea) respectively (17–18), and it traces to the last universal common ancestor (19–20).

The most common transition metal in enzymes is Fe, which usually occurs in electron-transferring FeS clusters (21). As an example, formyl-methanofuran dehydrogenase of the acetyl-CoA pathway in methanogens contains 46 electron-transferring 4Fe4S clusters (22). By contrast, Ni is rarely required by enzymes, and when it occurs, it participates in catalysis at the active site (reviewed by (23–24)). It is found in the active site of (i) urease—the first protein ever crystallized (*25*)—, of (ii) glyxoylase in methylglyoxal detoxification (26), of (iii) acireductone dioxygenase in the methionine salvage pathway) (27), and of (iv) Ni-superoxide dismutase (28–30), and in the active site of three enzymes of the acetyl-CoA pathway: In [Fe-Ni] hydrogenases (Hyd), Ni catalyzes the H_2_ dependent reduction of ferredoxin (31–36). In carbon monoxide dehydrogenase (CODH) Ni catalyzes the reduction of CO_2_ to CO (14, 34, 37–41). In acetyl-CoA synthase (ACS) Ni catalyzes the synthesis of acetyl-CoA from CO, CoASH and a methyl group (14, 34, 39, 41–42). In methanogens, nickel in F_430_ also catalyzes the final step of methane synthesis at methyl-CoM reductase (MCR) (43–47). The antiquity CO_2_ fixation via the acetyl-CoA pathway (13, 18) and the 3.5 Ga age of methanogenesis (48) trace Ni-based catalysis to the onset of biochemical evolution (49).

Although Ni in modern enzymes and cofactors is typically coordinated by sulfur or nitrogen atoms, (24, 50) this need not represent the ancestral state of catalytic Ni at origins. Inorganic NiS complexes can catalyze the ACS reaction starting from CO and methyl groups (51). The reduction of CO_2_ using NiS or FeS does not, however, take place unless external potentials of ∼1000 mV are applied (52–54), whereby that catalysis is afforded not by metal sulfides, but by native metals that are formed from the sulfides on the electrode during the electrochemical reaction (55). Without electrodes, pure Ni^0^ is an excellent catalyst of CO_2_ reduction. In the absence of enzymes, Ni^0^ in water catalyzes the H_2_-dependent reduction of CO_2_ to the products of the acetyl-CoA pathway—formate, acetate, and pyruvate—in hours to days at 50–100°C (56). That is, Ni^0^ nanoparticles replace the function of 10 enzymes and 10 cofactors of the acetyl-CoA pathway, which require the activity of 127 enzymes in cells (57). FeNi alloys and FeCo alloys as well as Fe^0^ alone catalyze the same spectrum of reactions as Ni^0^ (59–63). Beyond the reactions of the acetyl-CoA pathway, Ni^0^ catalyzes reactions of the reverse TCA cycle and H_2_-dependent reductive aminations of various 2-oxoacids to amino acids (64). Ni^0^ will catalyze the reduction of NADH (*E*_0_’ = –320 mV) with H_2_ (*E*_0_’ = –414 mV) (65–66), and it will reductively aminate pyridoxal to pyridoxamine using H_2_ (67), though it will not reduce low potential ferredoxin (*E*_0_’ = –450 mV) with H_2_, whereas Fe^0^ (*E*_0_ = –440 mV at pH0; *E*_0_ ca. –800 mV at pH14 (68) will (69).

Native nickel and its alloys are naturally deposited in serpentinizing hydrothermal vents by reduction of divalent metal ions with H_2_ generated during the serpentinization process (70–72). Nickel alloys are implicated in the purely geochemical synthesis of methane (73) that occurs in modern serpentinizing hydrothermal systems (74). Given that serpentinization has been going on since there was liquid water on Earth (3), nickel-based CO_2_ reduction and aminations likely operated before enzymes ever existed and continued to operate during the earliest phases of biochemical evolution as the first enzymes and pathways were arising (64, 66–67, 69, 75–76).

While the utility of Ni in enzymes lies in its ability to readily undergo changes of valence state (9, 77), its utility in catalysis of H_2_-dependent reductions of organic compounds lies in its ability to absorb and activate H_2_—a property that has been exploited in organic chemistry for over 100 years (78). Nickel particles avidly bind H_2_ as nickel hydride, Ni–H, at particle surfaces. At 1 atm H_2_, essentially all Ni atoms exposed at the surface of a Ni^0^ nanoparticle are occupied as Ni–H hydride (79). At higher H_2_ partial pressures, H diffuses into deeper atom layers as well, and at 30 atm of H_2_, a 2.7 nm diameter Ni nanoparticle can bind up to 3% H_2_ by weight (78). For comparison to natural and biological systems, the effluent of serpentinizing hydrothermal vents contains 1–10 atm H_2_ (80), methanogens require only 10^−5^–10^−4^ atm H_2_ to grow (81), while acetogens require 6·10^−5^–10^−3^ atm H_2_, depending on the strain (82). Ni–H formation from Ni^0^ and H_2_ is both spontaneous and facile (79), yet it is not a redox reaction, as both Ni and H remain in their elemental state. Like Co (83–84), Mo (85–86), and Fe (87–88) nickel also occurs in cofactors: the Ni-tetrapyrrole F_430_ of methanogens and in the nickel-pincer nucleotide (NPN) cofactor (89) first described in lactate racemase (90.).

The midpoint potential of Ni^0^ oxidation to Ni^2+^ under physiological conditions is *E*_0_’ = –260 mV, sufficient to reduce a number of common biological substrates, for example pyruvate to lactate (*E*_0_’ = –190 mV) in the absence of H_2_. With rare exceptions (67), the use of Ni^0^ in experimental reconstructions of metabolic origin so far has focused on H_2_-dependent reductions. Here we investigate reactions relevant to prebiotic chemistry using Ni^0^ as catalyst and reductant.

## RESULTS

In the presence of H_2_, nickel supplied as commercial nickel silicate powder (Ni-SiO_2_/Al_2_O_3_) catalyzes the reduction of pyruvate to lactate and, when ammonium is added in addition, pyruvate is reductively aminated to alanine (64, 67). However, the same reactions take place, though to a lesser extent, in the absence of H_2_. We incubated 20 mM pyruvate with and without 20 mM ammonium chloride in the presence or absence of Ni-SiO_2_/Al_2_O_3_ catalyst under 5 bar Ar. After 18h at 100°C and pH 11, a pH typical of serpentinizing hydrothermal systems (80, 91), 42% of pyruvate was reduced to lactate in the presence of Ni-SiO_2_/Al_2_O_3_ and 6.4% of pyruvate was reductively aminated to alanine in the presence of Ni-SiO_2_/Al_2_O_3_ and ammonia **(Figure 1).** Without the catalyst, no alanine or lactate accumulates. When Ni^0^ is present but ammonia is absent, only lactate is observed at a conversion rate of 49% (**Figure 1**)

**Figure 1.**
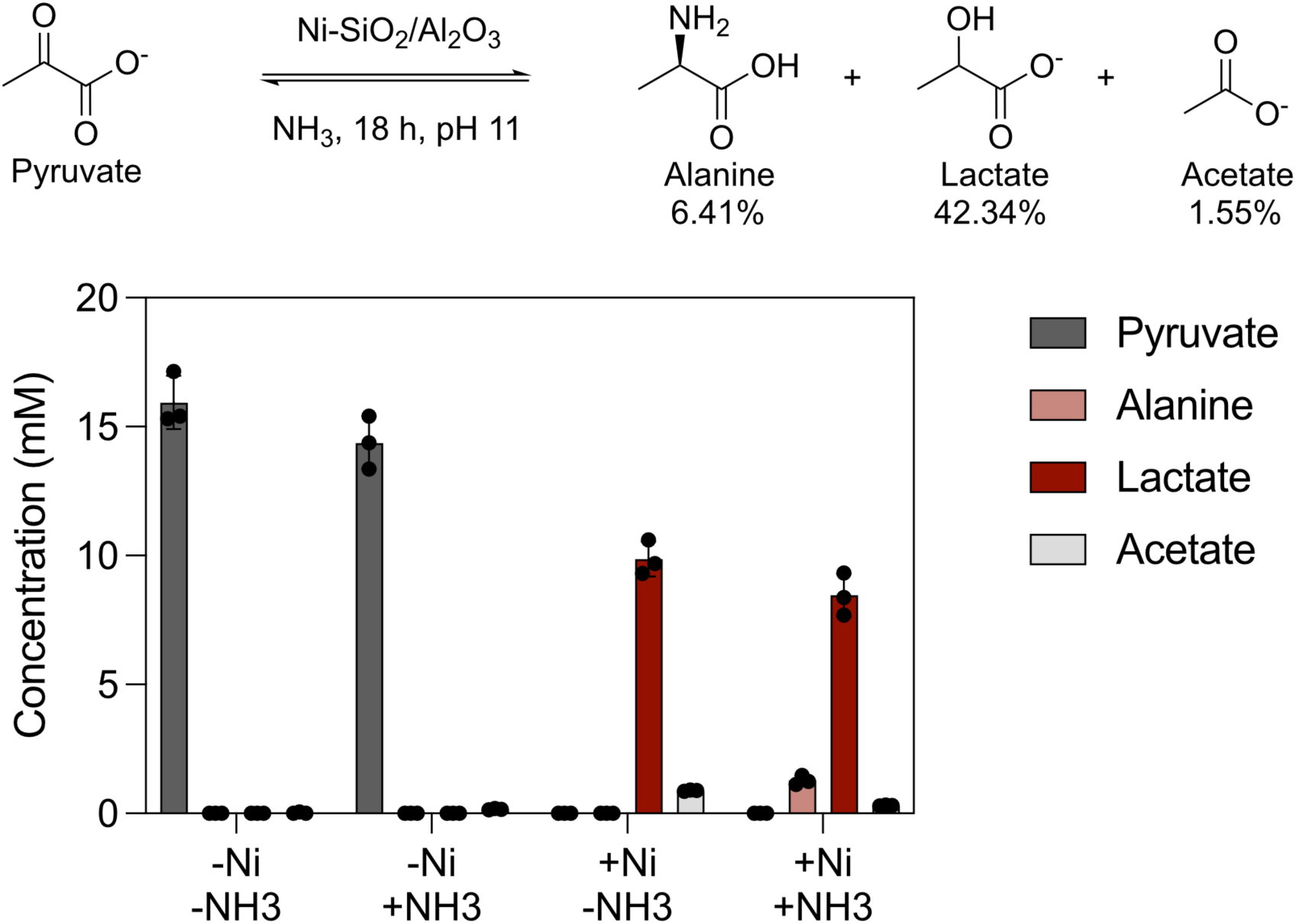
Alanine synthesis in the presence of nickel catalyst. Initial concentrations were 20 mM pyruvate and 200 mM ammonium chloride. Ni-SiO_2_/Al_2_O_3_ (1 mmol of metal atoms) was added as solid phase powder in a total reaction volume of 1.5 mL. The reaction was performed under a 5 bar Ar atmosphere, initial pH 11 with KOH, the reaction time was 18 h at 100°C. No H_2_ was added. Error bars in the figure indicate standard deviation (SD). Reactions were performed in triplicates.

This suggested that under the conditions of actively serpentinizing vents, nickel (*E*_0_’ = –260 mV), present as 1 mmol metal atoms versus 30 mmol pyruvate in the reaction, is acting not only a catalyst, but also as an electron donor for lactate synthesis (*E*_0_’ = –190 mV) as well. However, pyruvate could also be undergoing oxidation during the reaction and thus could, in principle, also serve as an electron donor. To test this, we examined parameters impacting the reduction of pyruvate to lactate.

First we examined the effect of pH. In actively serpentinizing hydrothermal systems, pH can typically vary in a range of 8-10, reaching pH 12 or higher in hyperalkaline systems (Cedars, (92); Oman ophiolite, (93); Hakuba happo, (94)). After 2 hours at 100°C at pH 7–11, we observed 86% conversion of pyruvate to lactate at pH 9 (**Figure 2**), such that pyruvate cannot be serving as the (sole) electron source. Acetate formation was very low, and increases with pH, whereas lactate peaks at pH 9.

**Figure 2.**
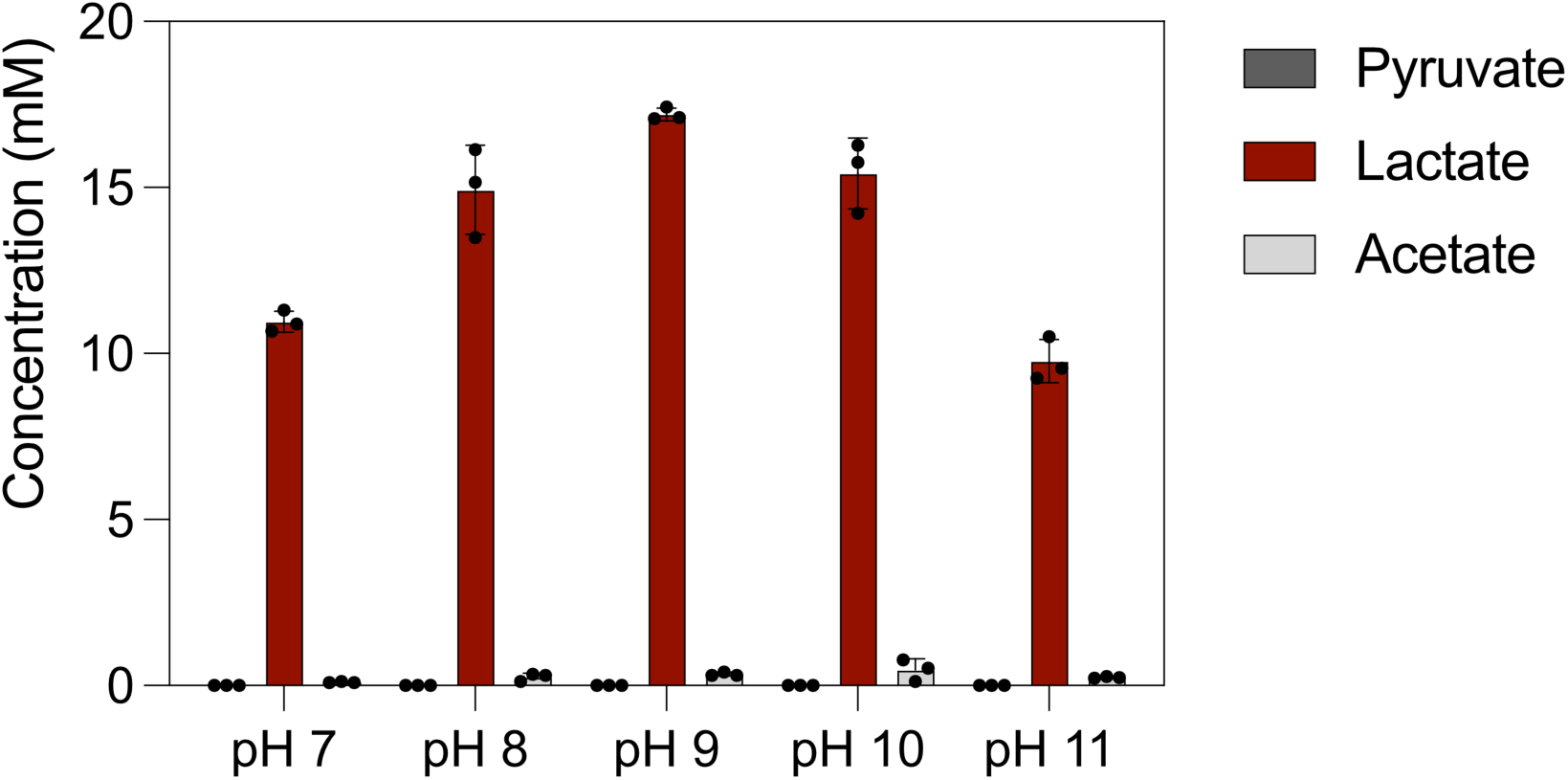
Effect of pH on lactate synthesis in the presence of nickel catalyst. The pH was set to 7, 8, 9, 10, and 11 with KOH, respectively, and the reaction time was set to 2 h. Other reaction parameters are as in Figure 1.

Temperature gradients exist in serpentinizing hydrothermal vents. Uncatalyzed biological reactions generally proceed more rapidly at increased temperature (5). Pyruvate reduction did not proceed at temperatures below 60°C and was essentially complete after 2 hours (**Figure 3**) at 80–100°C with 82-86% conversion. In 18 hour reactions, lactate decreases, likely due to thermal sensitivity (95).

**Figure 3.**
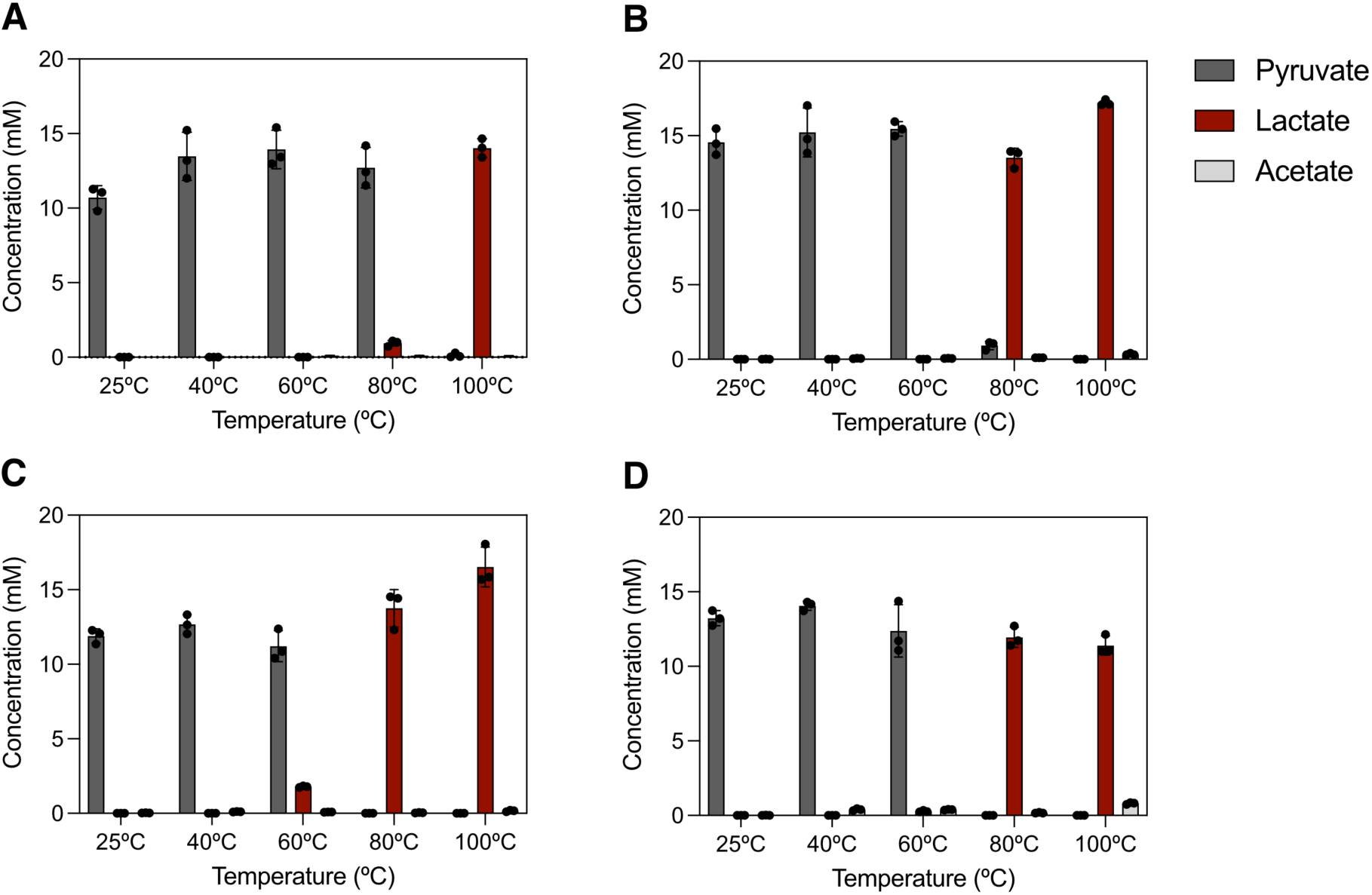
Effect of temperature and time on lactate synthesis in the presence of nickel catalyst. Parameters are as in Figure 1 unless otherwise indicated. The reaction time was 1h **(A)**, 2h **(B)**, 4h **(C)**, and 18 h **(D)**, respectively. Temperature was 25°C, 40°C, 60°C, 80°C, and 100°C. The pH was set to 9 with KOH.

Using commercial nano nickel powder and micro nickel powder, only partial reduction to lactate was observed after 18 hours (**Supplemental Figure 1**) with a yield of 37% with nano nickel powder, and 15.9% with micro nickel powder, but no reduction was observed after 2 hours. The silica matrix itself does not reduce pyruvate (**Supplemental Figure 1**).

Pyruvate was reduced to lactate using as little as 0.05 mmol of Ni per reaction, and the yield steadily increases when to 0.66 mmol of Ni per reaction. The yield plateaus at 1 mmol of Ni as Ni-SiO_2_/Al_2_O_3_ with 86% conversion (**Figure 4**).

**Figure 4.**
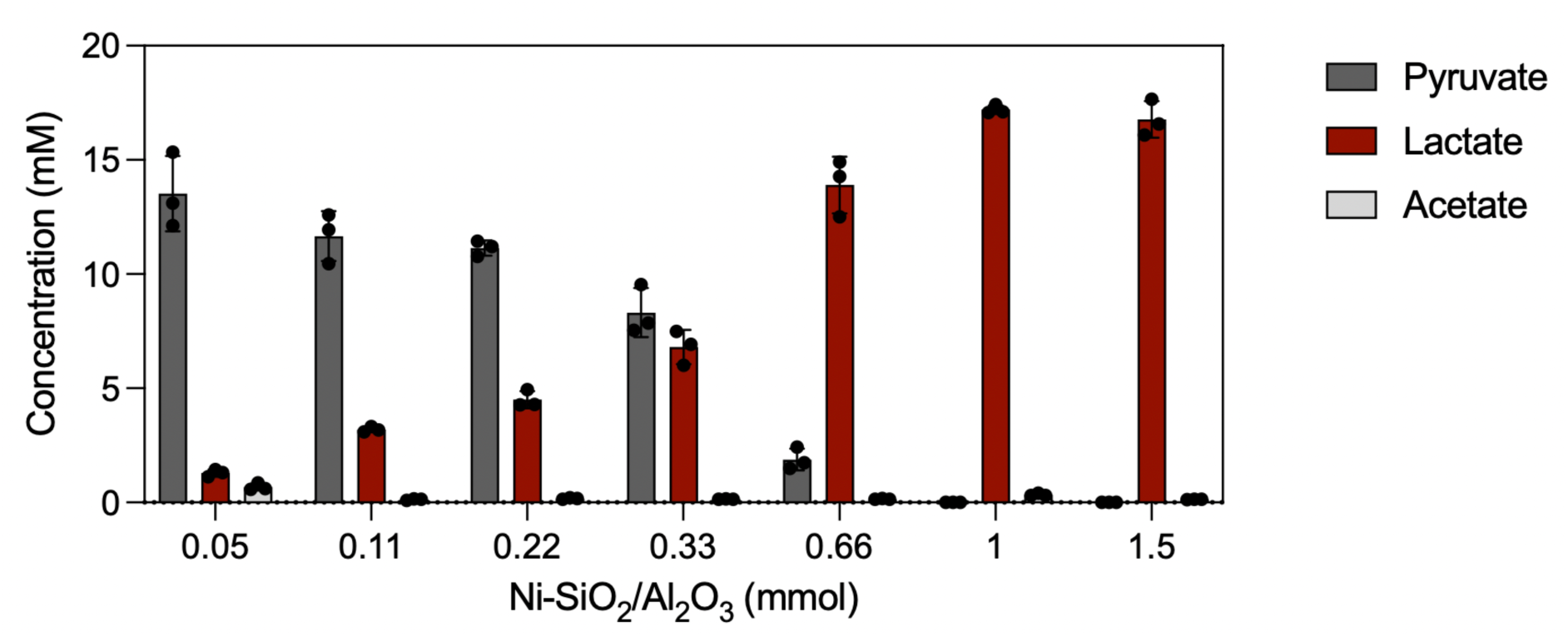
Effect of catalyst concentration on lactate synthesis in the presence of nickel catalyst. Parameters are as in Figure 1 unless otherwise indicated. Ni-SiO_2_/Al_2_O_3_ was added as solid phase at a concentration of 0.05, 0.11, 0.22, 0.33, 0.66, 1, and 1.5 mmol, respectively. The pH was 9 (KOH), and reaction time was 2 h. Error bars in the figure represent standard deviation (SD). Each reaction was performed in triplicates.

Kaur et al. (2024) converted 2-oxoglutarate, 4-methyl-2-oxopentaonate, and 3-methyl-2-oxopentaonate into the corresponding amino acids using Ni/H_2_ and NH_3_. Using 20 mM 2-oxoacid and 1.5 mM Ni as Ni-SiO_2_/Al_2_O_3_ under 5 bar Ar at pH 9 and 100°C, the reduction of 2-oxoglutarate to 2-hydroxyglutarate (*E*_0_’ = –337 mV) had a 100% conversion rate after 2h, while the reduction of 4-methyl-2-oxopentanoate to hydroxyisocaproate (*E*_0_’ = –344 mV) was 97% complete and the reduction of 3-methyl-2-oxopentanoate to 2-hydroxy-3-methylvalerate (*E*_0_’ = –357 mV) underwent 64% conversion (**Figure 5 A-C**). We also tested the possibility of nickel performing double bond reductions. Under the optimized conditions, fumarate was reduced to succinate (*E*_0_’ = –130 mV) with a 100% conversion rate (**Figure 5 D**)

**Figure 5.**
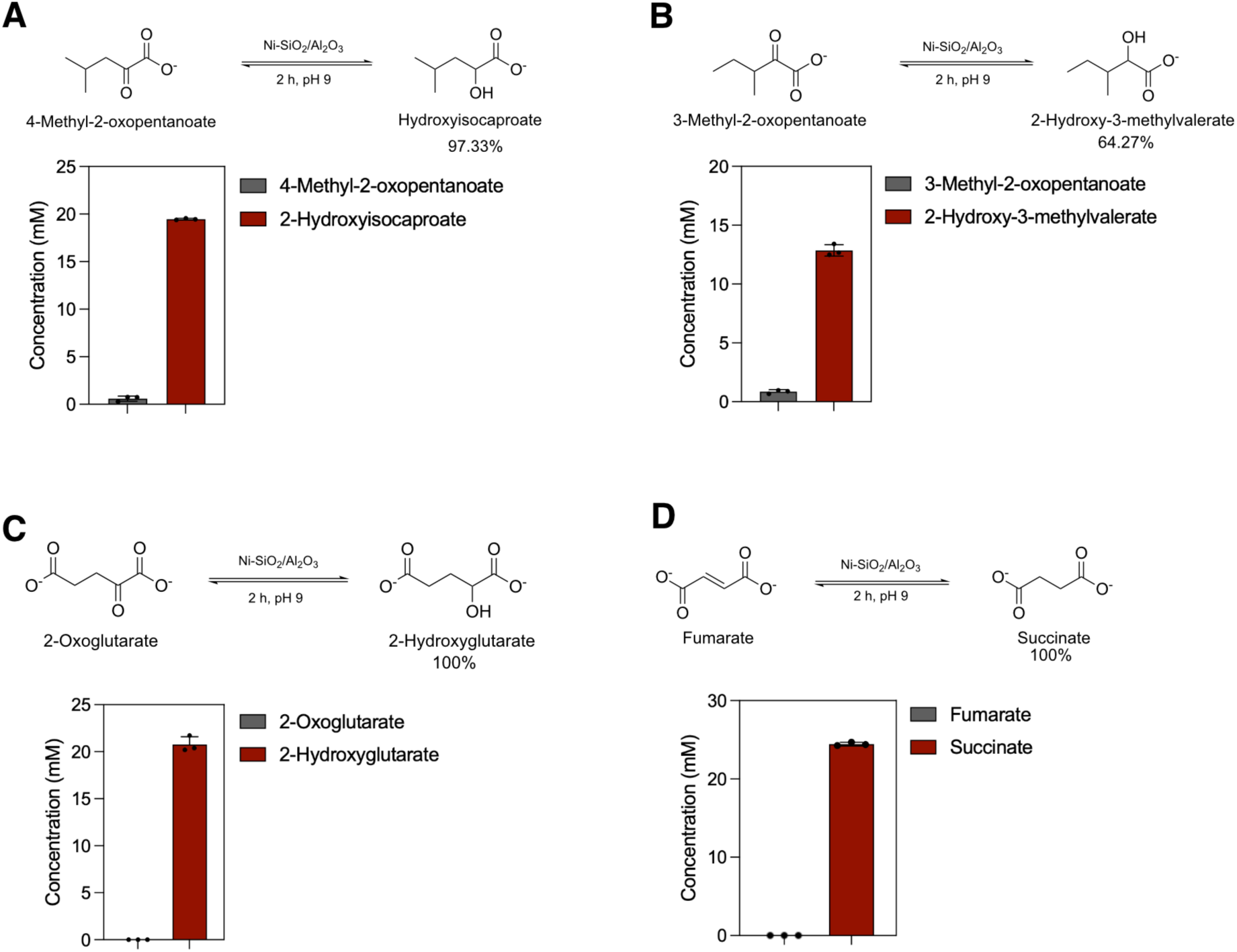
Ketone and double bond reduction in the presence of nickel catalyst. Parameters are as in Figure 1 unless otherwise indicated. Educt concentration was set to 20 mM. The reaction time was set to 2 h, and pH was set to 9 with KOH **A.** 2-Hydroxyisocaproate synthesis from 4-methyl-2-oxopentanoate. **B.** 2-Hydroxy-3-methylvalerate synthesis from 3-methyl-2-oxopentanoate. **C.** 2-Hydroxyglutarate synthesis from 2-oxoglutarate. **D.** Succinate synthesis from fumarate.

We investigated the reductive amination of these alpha-ketoacids to their corresponding amino acids using nickel as the reductant, adjusting the length of the reaction time (72h) and the pH (pH 11) to improve yields. Under these conditions, nickel has a lower midpoint potential (96), allowing for the reductive amination of the respective compounds. We observed that 4-methyl-2-oxopentanoate underwent reductive amination to leucine (*E*_0_’ = –386 mV) with 6.6% conversion rate and 3-methyl-2-oxopentanoate was reductively aminated to isoleucine (*E*_0_’ = –419 mV) with a 4.8% conversion rate (**Figure 6 A-B**). Reductive amination of 2-oxoglutarate did not generate glutamate (*E*_0_’ = –380 mV) but its cyclic peptide derivative 5-oxoproline, which is known to occur at high temperature and high pH (97). The formation of 5-oxoproline indicates reductive amination 2-oxoglutarate at 57% (**Figure 6 C**).

**Figure 6.**
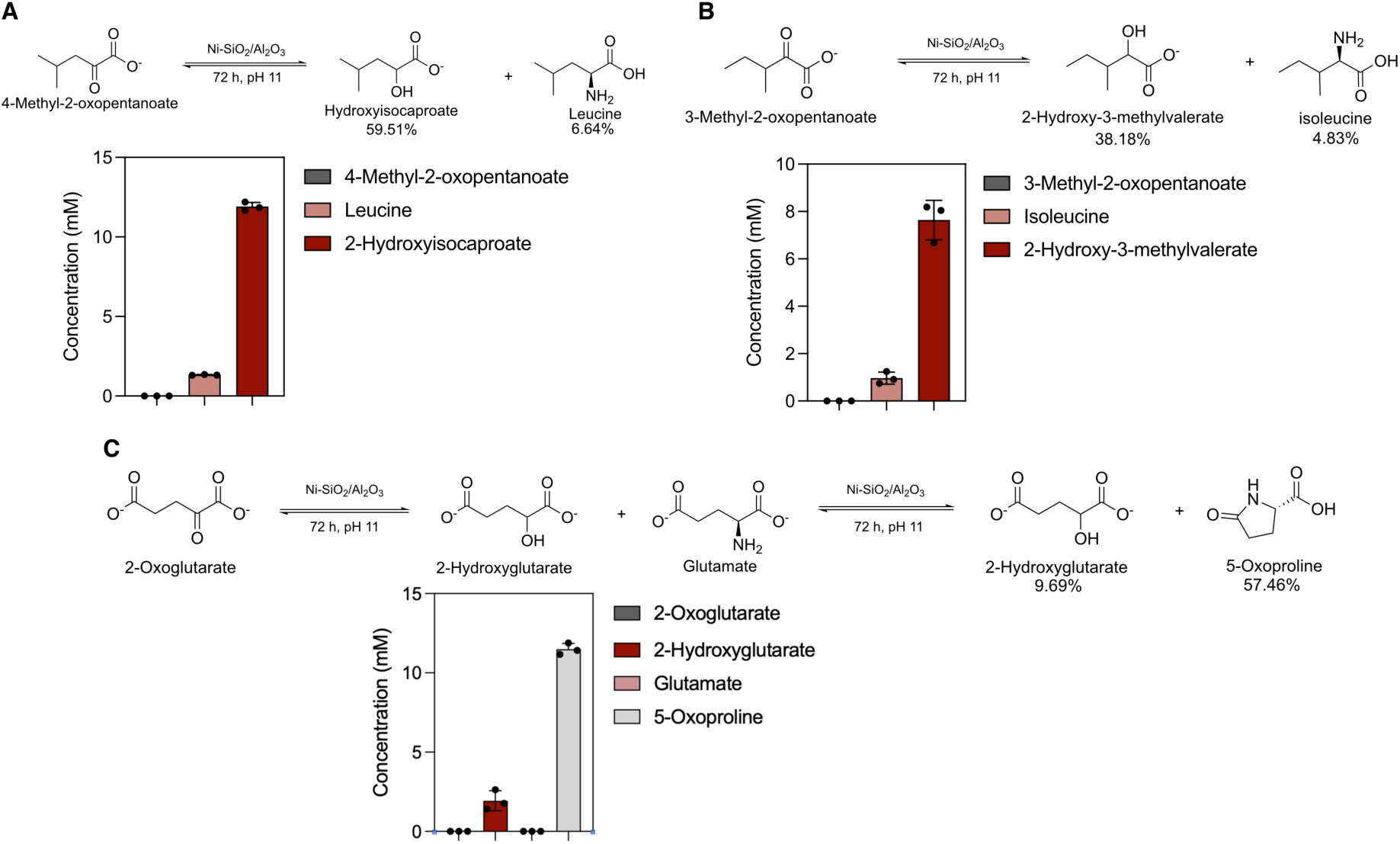
Reductive amination in the presence of nickel catalyst. Parameters are as in Figure 1 unless otherwise indicated. The reaction time was 72 h. **A.** Leucine synthesis from 4-methyl-2-oxopentanoate. **B.** Isoleucine synthesis from 3-methyl-2-oxopentanoate. **C.** 5-Oxoproline synthesis from 2-oxoglutarate.

## Discussion

Serpentinizing hydrothermal systems are interesting sites for the origin of metabolism because they generate a constant supply of H_2_ for CO_2_ reduction (49, 98–99) and because there is broad congruence between reactions catalyzed under hydrothermal conditions and reactions of metabolism (19, 97). The process of serpentinization furthermore generates highly alkaline effluent, pH 9-12, producing strongly reducing conditions with potentials on the order of –800 mV or more (90), which are sufficient to reduce divalent metals to native metals (70, 100–104). Native metals deposited in serpentinizing hydrothermal vents include Fe, Co, Ni, Pd, other platinum group elements (PGE) and their alloys (70, 104–106). These metals activate H_2_ via chemisorbtion and to catalyze organic reactions (60–63) under conditions conducive to the origin of metabolism. A number of recent studies have shown that Ni^0^ can promote the conversion of CO_2_ to organic acids using H_2_ as the reductant (60, 62–63), retracing rather exactly the reactions of the acetyl CoA pathway (60, 62–63) and reactions of the reverse TCA cycle (58, 64, 107). In water, Ni^0^, Fe^0^ and Co^0^ are furthermore compatible with cofactors, catalyzing the H_2_ dependent reduction of NAD^+^ (65–66), while Ni^0^ efficiently catalyzes the H_2_ dependent reduction of pyridoxal (67), and Fe^0^ catalyzes the reduction of 4Fe4S clusters in ferredoxin (69). With a midpoint potential of 440 mV, Fe^0^ can, and does, generate H_2_ in water, and can reduce CO_2_ without additional reductants (60). But Ni^0^ (*E*_0_’ = –270 mV) is a milder reductant.

Here we have shown that under serpentinizing conditions (high pH, temperature 50-100 °C) Ni^0^ can serve as a catalyst and reductant for reduction of 2-oxo groups to hydroxyl, reduction of double bonds, and reductive amination of 2-oxo acids to amino acids. These findings underscore the broad compatibility of native Ni with metabolic reactions. In modern metabolism, Ni is usually coordinated by S, C and N ligands in enzymes and cofactors as the divalent ion, but can undergo valance state changes during enzymatic reactions (23–24, 45). The extremely broad range of biochemical reactions that Ni^0^ can catalyze with or without H_2_ as a reductant and their similarity, often identity, to the reactions of metabolism in terms of reactants and products indicate the Ni^0^ was involved in metabolic origin. This is outlined in **Figure 7**.

**Figure 7.**
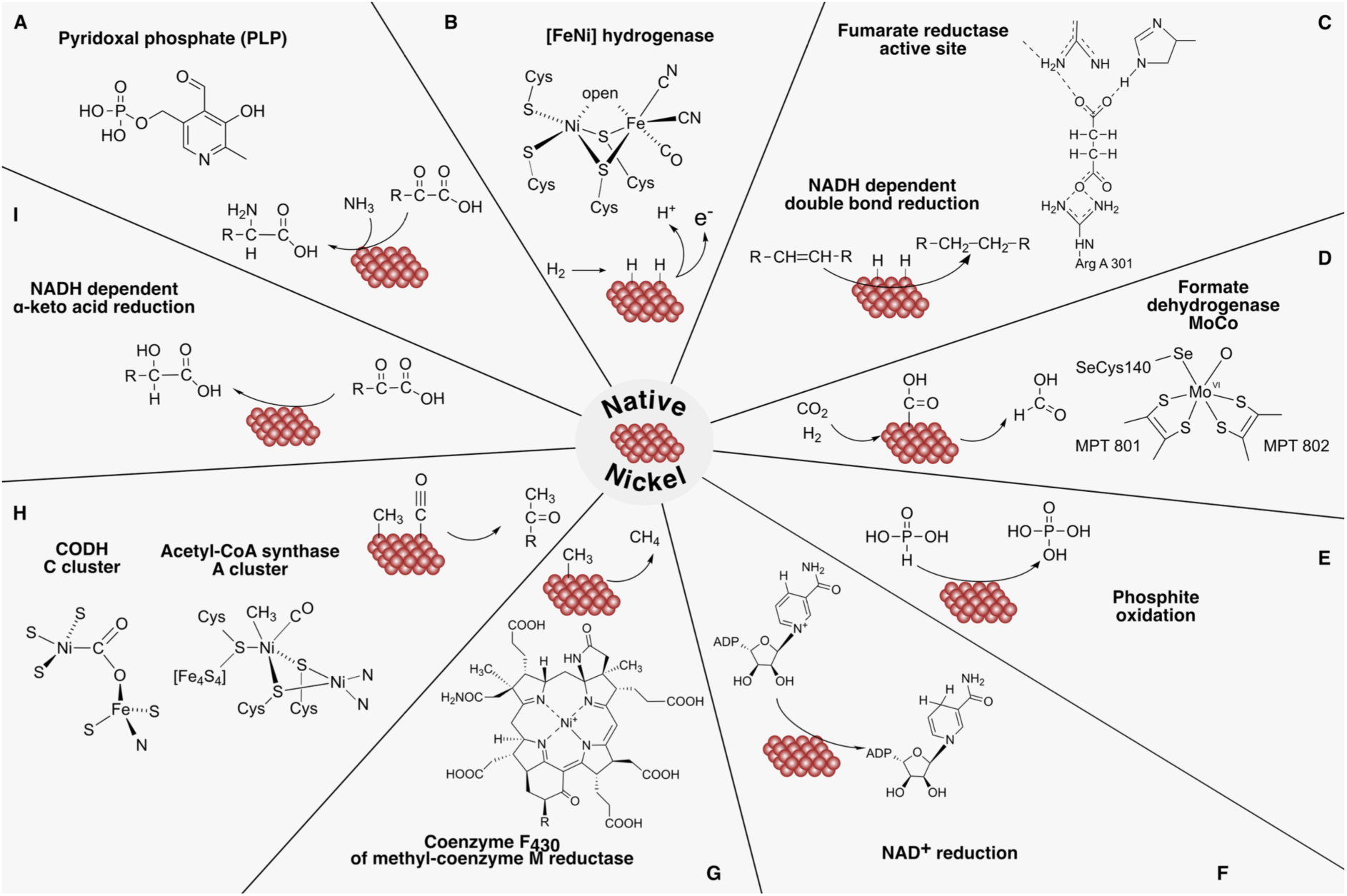
Broad substrate specificity of nickel as a primordial catalyst in metabolism. **(A)** Reductive amination: the presence of pyridoxal improves the yield (67), but the reaction also proceeds in the presence of Ni^0^ alone (this paper). **(B)** H_2_ oxidation: structure of carbon-Ni bond present in [FeNi] hydrogenase (31–32). Ni^0^ activates H_2_ (60) **(C)** Double bond reduction: active site of fumarase taken from (108), the presence of Ni^0^ alone catalyzes the reaction (this paper). **(D)** CO_2_ fixation: structure of molybdenum cofactor of formate dehydrogenase (109); Ni^0^ catalyzes the reaction under hydrothermal vent conditions (60). **(E)** Phosphite oxidation (97). **(F)** NAD^+^ reduction (65–66). **(G)** Methane formation: Ni^0^ generates methane from H_2_ and CO_2_ (76), the biological reaction requires coenzyme F_430_ in methyl-coenzyme M reductase (43). **(H)** CO_2_ fixation: structure of the carbon-Ni bonds present in carbon monoxide dehydrogenase (37–40), and acetyl-CoA synthase A-cluster (39–42); Ni^0^ catalyzes acetyl synthesis under hydrothermal vent conditions (60). **(I)** α-keto acid reduction: catalyzed by nickel (this paper).

Yet if Ni^0^ is so effective as a catalyst, one might ask why Ni^0^ is not used as a catalyst by enzymes or cofactors in modern metabolism. Cells can easily generate the potential needed to convert Ni^2+^ to Ni^0^, with the help of low potential reduced ferredoxin (*E*_o_ = –450 mV) (110) or using H_2_ with electron bifurcation (111).

Why is Ni^0^ not used by metabolism in cells? The answer, we suggest, is specificity. Broad substrate specificity is a classical trait of ancient enzymes (112–113). The first enzymes likely had very broad substrate specificity for a given kind of reaction (reductive amination for example) and diversified into ancient enzyme families each with greater substrate specificity, so that the reactions of (proto-) metabolism proceeded along orderly and well defined lines. Native nickel catalyzes too many different kinds of reactions (**Figure 7**). Ni^0^ catalyzes H_2_ activation (60, 62, 67) as in hydrogenases (32–36) and CO_2_ reduction to formate, acetate and pyruvate (60, 62) including the formation of C–C bonds (60, 62–64) as in the acetyl CoA pathway (18, 60). It catalyzes reactions of the rTCA cycle (58, 64, 107). And, as shown here, Ni^0^ catalyzes keto to alcohol reductions, reductive transaminations, and the reduction of double bonds in the absence of additional reductants. All of those reactions take place in water across a broad neutral to alkaline pH range and a wide range of biologically relevant temperatures. The extremely broad substrate specificity of Ni^0^ would make it an uncontrolled catalyst in a regulated metabolism, as cells could not turn it off. In a cytosol full with hundreds of metabolites present in µM to mM concentrations, Ni^0^ would have no choice but to react with whatever comes along first. The utility of Ni^0^ in prebiotic biochemical synthesis—capable of performing dozens of different reactions with CO_2_, NH_3_, H_2_ and organic moieties—is its liability in regulated metabolism. The catalytic versatility of Ni^0^ was useful in generating organic molecules at origins, but detrimental as enzymatic metabolism reached a state resembling that of modern cells. In order for natural selection in biochemical evolution to take place, enzymes had to be in control of the reactions they catalyzed. That was possible with Ni^2+^ coordinated by N, S, and C in enzymes and cofactors (24) (**Figure 7**), but not with Ni^0^, which is likely why it was left behind when metabolism escaped the hydrothermal environment (49, 97) within which it arose.

## MATERIAL AND METHODS

### Reaction conditions

Each reaction contained 20 mM of pyruvate (Sigma-Aldrich), or of its respective educt (2-oxoglutarate, 3-methyl-2-oxopentanoate, 4-methyl-2-oxopentanoate, fumarate; Sigma-Aldrich); and ammonium chloride when needed, dissolved in HPLC grade water (VWR International, Germany). The pH was adjusted from 7 to 11, depending on the experimental conditions required, by adding 1 M KOH. Ni_2_-Al_2_/O_3_ (Sigma-Aldrich) was used as a catalyst, added in a concentration between 0.03 mM to 1 mM, depending on the experimental conditions required, per mL of reaction volume, with a final reaction volume of 1.5 mL. Samples were prepared in glass vials (5 mL, Rotilabo-Rollrändfläschen ND20, Roth) and closed with metal lids (VWR International, Germany). Lids were punctured before being placed inside the reactor (Berghoff BR-300 with BTC-3000 temperature controller) to allow for gas exchange. Reactors were filled with 5-bar argon (99.996%; Messer, Switzerland). Reactions were performed at 25, 40, 60, 80 or 100°C over 1, 2, 4, 18 or 72 hours while stirring at 650 rpm.

When the reactions were over, reactors were depressurized and the reaction contents were transferred to 2 mL Eppendorf tubes, and centrifuged for 15 min at 13.000 rpm (Biofuge Fresco, Heraeus) to separate the reaction phases (metal contents and supernatant). The supernatants were then analyzed by NMR.

### Product identification

To prepare the samples for NMR analysis, 600 µL of each sample were transferred into NMR tubes (VWR International, Germany). As a reference for calibration, DSS (2,2-dimethyl-2-silapentane-5-sulfonate) was added, reaching a final concentration of 1 mM. NMR spectra were measured on a Bruker Avance III – 600 MHz spectrometer by the Center for Molecular and Structural Analytics at Heinrich Heine University Düsseldorf. Spectra were analyzed using Chenomx NMR Suite Version 9.02 software.

### Data availability

The data that supports the findings of this study are available in [Figure 1 -6] and the supplementary material of this article. Any raw data not specifically shown in the paper will be available upon request.

## Acknowledgments

This project has received funding from the European Research Council (ERC) under the European Union’s Horizon 2020 research and innovation program (grant agreement no. 101018894). For funding, W.F.M. thanks the ERC (101018894), the Deutsche Forschungsgemeinschaft (MA 1426/21-1) and the Volkswagen Foundation (Grant 96_742). We thank the Center for Molecular and Structural Analytics, Heinrich Heine University (CeMSA@HHU) for recording the NMR-spectroscopic data. We thank Joseph Moran, Harun Tüysüz, and Mirko Basen for many helpful discussions.

## Author contributions

C.G.G., M.B., and W.F.M. designed the research. C.G.G. performed the experiments. C.G.G., M.B., and W.F.M. interpreted and analyzed the data. C.G.G. and W.F.M. visualized the data. C.G.G., M.B. and W.F.M. wrote the paper. C.G.G., M.B., and W.F.M. edited the paper.

## REFERENCES

1. Lipmann, F. (1965) Projecting backward from the present stage of evolution of biosynthesis. In The Origins of Prebiological Systems and of their Molecular Matrices (Fox, S. W. ed), pp 259–280, Academic Press, New York, NY

2. Baross, J. A., and Hoffman, S. A. (1985) Submarine hydrothermal vents and associated gradient environments as sites for the origin and evolution of life. Orig. Life Evol. Biosph. 15, 327–345

3. Sleep, N. H., Bird, D. K., and Pope, E. C. (2011) Serpentine and the dawn of life. Philos. Trans. R. Soc. B. 366, 2857–2869

4. Stüeken, E. E., Buick, R., Guy, B. M., and Koehler, M. C. (2015) Isotopic evidence for biological nitrogen fixation by molybdenum-nitrogenase from 3.2 Gyr. Nature. 520, 666–669

5. Wolfenden, R., and Snider, M. J. (2001) The depth of chemical time and the power of enzymes as catalysts. Acc. Chem. Res. 34, 938–945

6. Eakin, R. E. (1963) An approach to the evolution of metabolism. Proc. Natl. Acad. Sci. USA. 49, 360

7. Hu, Y., and Ribbe, M. W. (2016) Nitrogenases–A tale of carbon atom(s). Angew. Chem. Int. Ed. 55, 8216–8226

8. Szaleniec, M., and Heider, J. (2025) Obligately tungsten-dependent enzymes–catalytic mechanisms, models and applications. Biochemistry. 64, 2154–2172

9. Ragsdale, S. W., and Wood, H. G. (1991) Enzymology of the acetyl-CoA pathway of CO_2_ fixation. Crit. Rev. Biochem. Mol. Biol. 26, 261–300

10. Drennan, C. L., Doukov, T. I., and Ragsdale, S. W. (2004) The metalloclusters of carbon monoxide dehydrogenase/acetyl-CoA synthase: A story in pictures. J. Biol. Inorg. Chem. 9, 511–515

11. Svetlitchnaia, T., Svetlitchnyi, V., Meyer, O., and Dobbek, H. (2006) Structural insights into methyltransfer reactions of a corrinoid iron-sulfur protein involved in acetyl-CoA synthesis. Proc. Natl. Acad. Sci. USA. 103, 14331–14336

12. Shima, S., Pilak, O., Vogt, S., Schick, M., Stagni, M. S., Meyer-Klaucke, W., et al. (2008) The crystal structrure of [Fe]-hydrogenase reveals the geometry of the active site. Science. 321, 572–575

13. Fuchs, G. (2011) Alternative pathways of carbon dioxide fixation: Insights into the early evolution of life? Annu. Rev. Microbiol. 65, 631–658

14. Yin, M. D., Lemaire, O. N., Jiménez, J. G. R., Belhamri, M., Shevchenko, A., Hummer, G., et al., (2025) Conformational dynamics of a multienzyme complex in anaerobic carbon fixation. Science. 387, 498–504

15. Berg, I. A., Kockelkorn, D., Ramos-Vera, W. H., Say, R. F., Zarzycki, J., Hügler, M., et al. (2010) Autotrophic carbon fixation in archaea. Nat. Rev. Microbiol. 8, 447–460

16. Berg, I. A. (2011) Ecological aspects of the distribution of different autotrophic CO_2_ fixation pathways. Appl. Envirom. Microbiol. 77, 1925–1936

17. Fuchs, G., and Stupperich, E. (1985) Evolution of autotrophic CO_2_ fixation. In Evolution of Prokaryotes (Schleifer, K. H., and Stackebrandt, E. eds), pp 235–25, Academic Press, London

18. Martin, W. F. (2020) Older than genes: The acetyl CoA pathway and origins. Front. Microbiol. 11, 817

19. Weiss, M. C., Sousa, F. L., Mrnjavac, N., Neukirchen, S., Roettger, M., Nelson-Sathi, S., et al. (2016) The physiology and habitat of the last universal common ancestor. Nat. Microbiol. 1, 1–8

20. Moody, E. R. R., Álvarez-Carretero, S., Mahendrajah, T. A., Clark, J. W., Betts, H. C., Dombrowski, N., et al. (2024) The nature of the last universal common ancestor and its impact on the early Earth system. *Nat*. Ecol. Evol. 8, 1654–1666

21. Lovenberg, W., Buchanan, B. B., and Rabinowitz, J. C. (1963) Studies on the chemical nature of clostridial ferredoxin. J. Biol. Chem. 238, 3899–3913

22. Wagner, T., Ermler, U., and Shima, S. (2016) The methanogenic CO_2_ reducing-and-fixing enzyme is bifunctional and contains 46 [4Fe-4S] clusters. Science. 354, 114–117

23. Ragsdale, S. W. (2009) Nickel-based enzyme systems. J. Biol. Chem. 284, 18571–18575

24. Neubeck, A., and Kirschning, A. (2026) Nickel: Geochemistry, biochemistry and its role in chemical and biological evolution. Earth-Sci. Rev. 272, 105324

25. Sumner, J. B. (1926) The isolation and crystallization of the enzyme urease: preliminary paper. J. Biol. Chem. 69, 435–441

26. Clugston, S. L., Barnard, J. F. J., Kinarch, R., Miedema, D., Ruman, R., Daub, E., et al. (1998) Overproduction and characterization of a dimeric non-zinc glyoxalase I from *Escherichia coli*: Evidence for optimal activation by nickel ions. Biochemistry. 37, 8754–8763

27. Dai, Y., Wensink, P. C., and Abeles, R. H. (1999) One protein, two enzymes. J. Biol. Chem. 274, 1193–1195

28. Youn, H. D., Kim, E. J., Roe, J. H., Hah, Y. C., and Kang, S. O. (1996) A novel nickel-containing superoxide dismutase from *Streptomyces* spp. Biochem. J. 318, 889–896

29. Youn, H. D., Youn, H., Lee, J. W., Yim, Y. I., Lee, J. K., Hah, Y. C., et al. (1996) Unique isozymes of superoxide dismutase in *Streptomyces griseus*. Arch. Biochem. Biophys. 334, 341–348

30. Kim, F. J., Kim, H. P., Hah, Y. C., and Roe, J. H. (1996) Differential expression of superoxide dismutases containing Ni and Fe/Zn in *Streptomyces coelicolor*. Eur. J. Biochem. 241, 178–185

31. Volbeda, A., Charon, M. H., Piras, C., Hatchikian, E. C., Frey, M., and Fontecilla-Camps, J. C. (1995) Crystal structure of the nickel-iron hydrogenase from *Desulfovibrio gigas*. Nature. 373, 580–587

32. Armstrong, F. A., and Albracht, S. P. J. (2005) [NiFe]–hydrogenases: spectroscopic ad electrochemical definition of reactions and intermediates. Philos. Trans. A. Math. Phys. Eng. Sci. 363, 937–954

33. Thauer, R. K., Kaster, A. K., Goenrich, M., Schick, M., Hiromoto, T., and Shima, S. (2010) Hydrogenases from methanogenic archaea, nickel, a novel cofactor, and H_2_ storage. Annu. Rev. Biochem. 79, 507–536

34. Nitschke, W., McGlynn, S. E., Milner-white, E. J., and Russell, M. J. (2013) On the antiquity of metalloenzymes and their substrates in bioenergetics. Biochim. Biophys. Acta. 1827, 871–881

35. Ogata, H., Lubitz, W., and Higuchi, Y. (2016) Structure and function of [NiFe] hydrogenases. J. Biochem. 160, 251–258

36. Harrison, D. J., Lough, A. J., and Fekl, U. (2018) A new structural model for NiFe hydrogenases: An unsaturated analogue of a classic hydrogenase model leads to more enzymes-like Ni-Fe distance and interplanar fold. Acta Crystallogr. E. 74, 1222–1226

37. Dobbek, H., Svetlitchnyi, V., Gremer, L., Huber, R., and Meyer, O. (2001) Crystal structure of a carbon monoxide dehydrogenase reveals a [Ni-4-Fe-5S] cluster. Science. 293, 1281–1285

38. Jeoung, J. H., and Dobbek, H. (2007) Carbon dioxide activation at the Ni,Fe-cluster of anaerobic carbon monoxide dehydrogenase. Science. 318, 1461–1464

39. Can, M., Armstrong, F. A., and Ragsdale, S. W. (2014) Structure, function, and mechanism of the nickel metalloenzymes, CO dehydrogenase, and acetyl-CoA synthase. Chem. Rev. 114, 4149–4174

40. Can, M., Giles, L. J., Ragsdale, S. W., and Sarangi, R. (2017) X-ray absorption spectroscopy reveals an organometallic Ni-C Bond in the CO-treated form of acetyl-CoA synthase. Biochemistry. 56, 1248–1260

41. Biester, A., Grahame, D. A., and Drennan, C. L. (2024) Capturing a methanogenic carbon monoxide dehydrogenase/acetyl-CoA synthase complex via cryogenic electron microscopy. Proc. Natl. Acad. Sci. USA. 121, e2410995121

42. Svetlitchnyi, V., Dobbek, H., Neyer-Klaucke, W., Meins, T., Thiele, B., Römer, P., et al. (2004) A functional Ni-Ni-[4Fe-4S] cluster in the monomeric acetyl-CoA synthase from *Carboxydothermus hydrogenoformans*. Proc. Natl. Acad. Sci. USA. 101, 446–451

43. Rospert, S., Böcher, R., Albracht, S. P. J., and Thauer R. K. (1991) Methyl-coenzyme M reductase preparations with high specific activity from H_2_ preincubated cells of *Methanobacterium thermoautotrophicum*. FEBS Lett. 291, 371–375

44. Goubeaud, M., Schreiner, G., and Thauer, R. K. (1997) Purified methyl-coenzyme-M reductase is activated when the enzyme-bound coenzyme F_430_ is reduced to the nickel(I) oxidation state by titanium(III) citrate. Eur. J. Biochem. 243, 110–114

45. Goenrich, M., Mahlert, F., Duin, E. C., Bauer, C., Jaun, B., and Thauer, R. K. (2004) Probing the reactivity of Ni in the active site of methyl-coenzyme M reductase with substrate analogues. J. Biol. Inorg. Chem. 9, 691–705

46. Wongnate, T., Sliwa, D., Ginovska, B., Smith, D., Wolf, M. W., Lehnert, N., et al. (2016) The radical mechanism of biological methane synthesis by methylcoenzyme M reductase. Science. 352, 953–958

47. Thauer, R. K. (2019) Methyl (alkyl)-coenzyme m reductases: Nickel F-430-containing enzymes involved in anaerobic methane formation and in anaerobic oxidation of methane or of short chain alkanes. Biochemistry. 58, 5198–5220

48. Ueno, Y., Yamada, K., Yoshida, N., Maruyama, S., and Isozaki, Y. (2006) Evidence from fluid inclusions for microbial methanogenesis in the early Archaean era. Nature. 440, 516–519

49. Martin, W. F., Baross, J., Kelley, D., and Russell, M. J. (2008) Hydrothermal vents and the origin of life. Nat. Rev. Microbiol. 6, 805–814

50. Shima, S., Huang, G., Wagner, T., and Ermler, U. (2020) structural basis of hydrogenotrophic methanogenesis. Annu. Rev. Microbiol. 74, 713–733

51. Huber, C., and Wächtershäuser, G. (1997) Activated acetic acid by carbon fixation on (Fe,Ni)S under primordial conditions. Science. 276, 245–247

52. Roldan, A., Hollingsworth, N., Roffey, A., Islam, H. U., Goodall, J. B. M., Catlow, C. R. A., et al. (2015) Bio-inspired CO_2_ conversion by iron sulfide catalysts under sustainable conditions. Chem. Commun. 51, 7501–7504

53. Kitadai N., Nakamura, R., Yamamoto, M., Takai, K., Li, Y., Yamaguchi, A., et al. (2018) Geoelectrochemical CO production: Implications for the autotrophic origin of life. Sci. Adv. 4, eaao7265

54. Kitadai, N., Nakamura, R., Yamamoto, M., Okada, S., Takahagi, W., Nakano, Y., et al. (2021) Thioester synthesis through geoelectrochemical CO_2_ fixation on Ni sulfides. Commun. Chem. 4, 37

55. Kitadai, N., Nakamura, R., Yamamoto, M., Takai, K., Yoshida, N., and Oono, Y. (2019) Metals likely promoted protometabolism in early ocean alkaline hydrothermal systems. Sci. Adv. 5, eaav7848

56. Beyazay, T., Belthle, K. S., Farès, C., Preiner, M., Moran, J., Martin, W. F., et al. (2023) Ambient temperature CO_2_ fixation to pyruvate and subsequently to citramalate over iron and nickel nanoparticles. Nat. Commun. 14, 570

57. Mrnjavac, N., Wimmer, J. L. E., Brabender, M., Schwander, L., and Martin, W. F. (2023) The moon-forming impact and the autotrophic origin of life. ChemPlusChem 88, e202300270

58. Muchowska, K. B., Varma, S. J., Chevallot-Beroux, E., Lethuillier-Karl, L., Li, G., and Moran, J. (2017) Metals promote sequences of the reverse Krebs cycle. *Nat*. Ecol. Evol. 1, 1716–1721

59. Varma, S. J., Muchowska, K. B., Chatelain, P., and Moran, J. (2018) Native iron reduces CO_2_ to intermediates and end-products of the acetyl-CoA pathway. *Nat*. Ecol. Evol. 2, 1019–1024

60. Preiner, M., Igarashi, K., Muchowska, K. B., Yu, M., Varma, S. J., Kleinermanns, K., et al. (2020) A hydrogen-dependent geochemical analogue of primordial carbon and energy metabolism. *Nat*. Ecol. Evol. 4, 534–542

61. Belthe, K. S., Beyazay, T., Ochoa-Hernández, C., Miyazaki, R., Foppa, L., Martin, W. F., et al. (2022) Effects of silica modification (Mg, Al, Ca, Ti, and Zr) on supported cobalt catalysts for H_2_-dependent CO_2_ reduction to metabolic intermediates. J. Am. Chem. Soc. 144, 21232–21243

62. Beyazay, T., Ochoa-Hernández, C., Song, Y., Belthe, K. S., Martin, W. F., and Tüysüz, H. (2023) Influence of composition of nickel-iron nanoparticles for abiotic CO_2_ conversion to early prebiotic organics. Angew. Chem. Int. Ed. 62, e202218189

63. Belthe, K. S., Martin, W. F., and Tüysüz, H. (2024) Synergistic effects of silica-supported iron-cobalt catalysts for CO_2_ reduction to prebiotic organics. ChemCatChem. 16, e202301218

64. Kaur, H., Rauscher, S., Werner, E., Song, Y., Yi, J., Kazöne, W., et al. (2024) A prebiotic Krebs cycle analog generates amino acids with H_2_ and NH_3_ over nickel. Chem. 10, 1528–1540

65. Henriques Pereira, D. P., Leethaus, J., Beyazay, T., do Nascimiento Vieira, A., Kleinermanns, K., Tüysüz, H., et al. (2022) Role of geochemical protoenzymes (geozymes) in primordial metabolism: specific abiotic hydride transfer by metals to the biological redox cofactor NAD^+^. FEBS J. 289, 3148–3162

66. Henriques Pereira, D. P., Xie, X., Stewart, S. V., Subrati, Z., Beyazay, T., Paczia, N., et al. (2025) Reduction of NAD and NMN on mineral surfaces with H_2_ reveals a functional role for the AMP moiety in a prebiotic context. Commun. Chem. 8, 318

67. Schlikker, M. L., Brabender, M. Schwander, L., Garcia Garcia, C., Burmeister, M., et al. (2025) Conversion of pyridoxal to pyridoxamine with NH_3_ and H_2_ on nickel generates a protometabolic nitrogen shuttle under serpentinizing conditions. FEBS J. 292, 3041–3055

68. Bard, A. J., Parsons, R., and Jordan, J. (1985) Standard Potentials in Aqueous Solution, 1st Ed., Routledge, New York.

69. Brabender, M., Henriques Pereira, D. P., Mrnjavac, N., Schlikker, M. L., Kimura, Z. I., Sucharitakul, J., et al. (2024) Ferredoxin reduction by hydrogen with iron functions as an evolutionary precursor of flavin-based electron bifurcation. Proc. Natl. Acad. Sci. USA. 121, e2318969121

70. Chamberlain, J. A., McLeod, C. R., Traill, R. J., and Lachance, G. R. (1965) Native metals in the muskox intrusion. Can. J. Earth Sci. 2, 188–215

71. Russell, M. J., Hall, A. J., and Martin, W. F. (2010) Serpentinization as a source of energy at the origin of life. Geobiology. 8, 355–371

72. Schwarzenbach, E. M., Vrijmoed, J. C., Engelmann, J. M., Liesengang, M., Wiechert, U., Rohne R., et al. (2021) Sulfide dissolution and awaruite formation in continental serpentinization environments and its implications to supporting life. J. Geophys. Res. Solid Earth. 126, e2021JB021758

73. Horita, J., and Berndt, M. E. (1999) Abiogenic methane formation and isotopic fractionation under hydrothermal conditions. Science. 285, 1055–1057

74. Etiope, G., Sherwood Lollar, B. (2013) Abiotic methane on Earth. Rev. Geophys. 51, 276–299

75. Mrnjavac, N., Schwander, L., Brabender, M., and Martin, W. F. (2024) Chemical antiquity in metabolism. Acc. Chem. Res. 57, 2267–2278

76. Song, Y., and Tüysüz, H. (2024) CO_2_ Fixation to prebiotic intermediates over heterogeneous catalysts. Acc. Chem. Res. 57, 2038–2047

77. Diekert, G., and Thauer, R. K. (1980) The effect of nickel on carbon monoxide dehydrogenase formation in *Clostridium thermoaceticum* and *Clostridium formicoaceticum*. FEMS Microbiol. Lett. 7, 187–189

78. Adkins, H., and Cramer, H. I. (1930) The use of nickel as a catalyst for hydrogenation. J. Am. Chem. Soc. 52, 4349–4358

79. Rana, S., Masli, N., Monder, D. S., and Chatterjee, A. (2022) Hydriding pathway for Ni nanoparticles: Computational characterization provides insights into the nanoparticle size and facet effect on layer-by-layer subsurface hydride formation. Comput. Mater. Sci. 210, 111482

80. Schwander, L., Brabender, M., Mrnjavac, N., Wimmer, J. L. E., Preiner, M., and Martin, W. F. (2023) Serpentinization as the source of energy, electrons, organics, catalysts, nutrients and pH gradients for the origin of LUCA and life. Front. Microbiol. 14, 1257597

81. Thauer, R. K., Kaster, A. K., Seedorf, H., Buckel, W., and Hedderich, R. (2008) Methanogenic archaea: ecologically relevant differences in energy conservation. Nat. Rev. Microbiol. 6, 579–591

82. Munoz, L., and Philips, J. (2023) No acetogen is equal: Strongly different H_2_ thresholds reflect diverse bioenergetics in acetogenic bacteria. Environ. Microbiol. 25, 2032–2040

83. Hodgkin, D. M. C. (1965) The structure of the corrin nucleus from X-ray analysis. Proc. R. Soc. A. 288, 294–305

84. Marques, H. M., and Brown, K. L. (1999) The structure of cobalt corrinoids based on molecular mechanics and NOE-restrained molecular mechanics and dynamics simulations. Coord. Chem. Rev. 190-192, 127–153

85. Johnson, J. L., Hainline, B. E., and Rajagopalan, K. V. (1980) Characterization of the molybdenum cofactor of sulfide oxidase, xanthine oxidase, and nitrate reductase. Identification of a pteridine as a structural component. J. Biol. Chem. 255, 1783–1786

86. Johnson, J. L., and Rajagopalan, K. V. (1982) Structural and metabolic relationship between the molybdenum cofactor and urothione. Proc. Natl. Acad. Sci. USA. 79, 6856–6860

87. Buurman, G., Shima, S., and Thauer, R. K. (2000) The metal-free hydrogenase from methanogenic archaea: evidence for a bound cofactor. FEBS Lett. 485, 200–204

88. Shima, S., Lyon, E. J., Sordel-Klippert, M., Kauß, M., Kahnt, J., Thauer, R. K, et al. (2004) The Cofactor of the Iron-Sulfur Cluster Free Hydrogenase Hmd: Structure of the Light-Inactivation Product. Angew. Chem. Int. Ed. 43, 2547–2551

89. Gatreddi, S., Chatterjee, S., Turmo, A., Hu, J., and Hausinger, R. P. (2024) A structural view of nickel-pincer nucleotide cofactor-related biochemistry. Crit. Rev. Biochem. Mol. Biol. 59, 402–417

90. Desguin, B., Zhang, T., Soumillon, P., Hols, P., Hu, J., and Hausinger, R. P. (2015) A tethered niacin-derived pincer complex with a nickel-carbon bond in lactate racemase. Science 349, 66–69

91. Boyd, E. S., Amenabar, M. J., Poudel, S., and Templeton, A. S. (2020) Bioenergetic constraints on the origin of autotrophic metabolism. Philos. Trans. R. Soc. A. 379, 20190151

92. Suzuki, S., Ishii, S., Hoshino, T., Rietze, A., Tenney, A., Morrill, P. L., et al. (2017) Unusual metabolic diversity of hyperalkaliphilic microbial communities associated with subterranean serpentinization at The Cedars. ISME J. 11, 1584–1598

93. Colman, D. R., Kraus, E. A., Thieringer, P. H., Rempfert, K., Templeton, A. S., Spear, J. R., et al. (2022) Deep-branching acetogens in serpentinized subsurface fluids of Oman. Proc. Natl. Acad. Sci. USA. 119, e2206845119

94. Nobu, M. K., Nakai, R., Tamazawa, S., Mori, H., Toyoda, A., Ijiri, A., et al. (2023) Unique H_2_-utilizing lithotrophy in serpentinite-hosted systems. ISME J. 17, 95–104

95. Komesu, A., Marins Martínez, P. H., Lunelli, B. H., Olivieira, J., Wolf Maclel, M. R., and Maclel Filho R. (2017) Study of lactic acid thermal behavior using thermoanalytical techniques. J. Chem. 2017, 4149592

96. Beverskog, B., and Puigdomenech, I. (1997) Revised Pourbaix diagrams for nickel at 25–300 °C. Corros. Sci. 39, 969–980

97. Mrnjavac, N., Hoffmann, N. K., Schlikker, M. L., Burmeister, M., Schwander, L., Garcia Garcia, C., et al. (2025) Gradual assembly of metabolism at a phosphorylating hydrothermal vent. *arXiv*, arXiv2510:08410

98. Sleep, N. H., Meibom, A., Fridriksson, T., Coleman, R. G., and Bird, D. K. (2004) H_2_-rich fluids from serpentinization: geochemical and biotic implications. Proc. Natl. Acad. Sci. USA. 101, 12818–12823

99. McCollom, T. M., and Seewald, J. S. (2013) Serpentinites, hydrogen, and life. Elements. 9, 129–134

100. Sinton J. M. (1976) Compositional relationships of Fe-Ni alloy and coexisting phases in serpentinite, Red Mountain, New Zealand. Mineral. Mag. 40, 792–794

101. Lorand, J. P. (1987) Cu-Fe-Ni-S mineral assemblages in upper-mantle peridotites from the Table Mountain and Blow-Me-Down Mountain ophiolite massifs (Bay of Islands área, Newfoundland): Their relationships with fluids and silicate melts. Lithos. 20, 59–76

102. Klein, F., Bach, W., Jöns, N., McCollom, T. M., Moskowitz, B., and Berquó, T. (2009) Iron partitioning and hydrogen generation during serpentinization of abyssal peridotites from 15°N of the Mid-Atlantic Ridge. Geochim. Cosmochim. Acta. 73, 6868–6893

103. Schwarzenbach, E. M., Vrijmoed, J. C., Engelmann, J. M., Liesengang, M., Wiechert, U., Rohne, R., et al. (2021) Sulfide dissolution and awaruite formation in continental serpentinization environments and its implications to supporting life. J. Geophys. Res. B. 126, e2021JB021758

104. Tamblyn, R., and Hermann, J. (2023) Geological evidence for high H_2_ production from komatiites in the Archaean. Nat. Geosci. 16, 1194–1199

105. Lawley, C. J. M., Petts, D. C., Jackson, S. E., Zagorevski, A., Pearson, D. G., Kjarsgaard, B. A., et al. (2020) Precious metal mobility during serpentinization and breakdown of base metal sulphide. Lithos. 354**-**355, 105278

106. Kutyrev, A., Kamenetsky, V. S., Kontonikas-Charos, A., Savelyev, D. P., Yakich, T. Y., Belousov, I. A., et al. (2023) Behaviour of platinum-group elements during hydrous metamorphism: constraints from awaruite (Ni_3_Fe) mineralization. Lithosphere. 2023, lithosphere_2023_126

107 Rauscher, S. A., and Moran J. (2022) Hydrogen drives part of the reverse Krebs cycle under metal or meteorite catalysis. Angew. Chem. Int. Ed. 61, e202212932

108. Lancaster, C. R. D., Kröger, A., Auer, M., and Michel, H. (1999) Structure of fumarate reductase from Wolinella suxinogenes at 2.2 Å resolution. Nature. 402, 377–385

109. Boyington, J. C., Gladyshev, V. N., Khangulov, S. V., Stadtman, T. C., and Sun, P. D. (1997) Crystal structure of formate dehydrogenase H: Catalysis involving Mo, molybdopterin, selenocysteine, and an Fe4S4 cluster. Science. 275, 1305–1308

110. Buckel, W., and Thauer, R. K. (2013) Energy conservation via electron bifurcating ferredoxin reduction and proton/Na(+) translocating ferredoxin oxidation. Biochim. Biophys. Acta. 1827, 94–113

111. Kaster, A. K., Moll, J., Parey, K., and Thauer, R. K. (2011) Coupling of ferredoxin and heterodisulfide reduction via electron bifurcation in hydrogenotrophic methanogenic archaea. Proc. Natl. Acad. Sci. USA. 108, 2981–2986

112. Jensen, R. A. (1976) Enzyme recruitment in evolution of new function. Annu. Rev. Microbiol. 30, 409–425

113. Khersonsky, O., and Tawfik, D. S. (2010) Enzyme promiscuity: A mechanistic and evolutionary perspective. Annu. Rev. Microbiol. 79, 471–505

