## Supplemental Figures for "Prebiotic aqueous reactions catalyzed by native nickel without hydrogen"

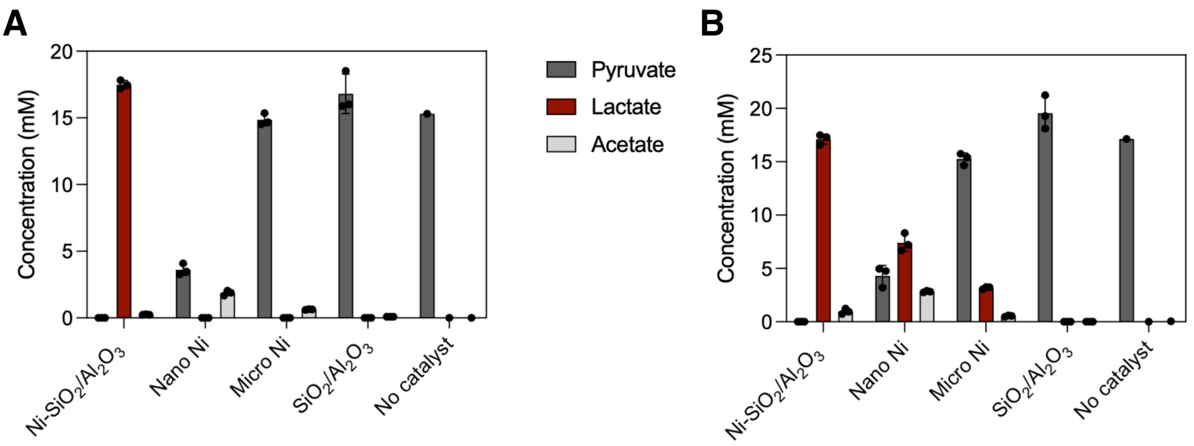


Supplemental Figure 1. **Effect of different catalysts on lactate synthesis.** Pyruvate concentration was set to 20 mM. The catalysts (Ni-$SiO_2$/$Al_2O_3$, Nano nickel powder, micro nickel powder, $SiO_2$/$Al_2O_3$) were added as 1.5 mmol of undissolved solid phase powder in a total reaction volume of 1.5 mL. The reaction was performed at 100°C, under a 5 bar argon atmosphere, pH was set to 9 with KOH. The reaction time was set to 2 **(A)**, and 18 **(B)**. Error bars in the figure represent standard deviation (SD). Each reaction was performed in triplicates.

| | Pyruvate [-$Ni^0$] [-$NH_3$] (mM) | Pyruvate [-$Ni^0$] [+$NH_3$] (mM) | Pyruvate [+$Ni^0$] [-$NH_3$] (mM) | Pyruvate [+$Ni^0$] [+$NH_3$] (mM) |
|---|---|---|---|---|
| [1] | 15.2761 | 15.4888 | n.d. | n.d. |
| [2] | 14.4107 | 13.3275 | n.d. | n.d. |
| [3] | 17.0135 | 14.3597 | n.d. | n.d. |

| | Alanine [-$Ni^0$] [-$NH_3$] (mM) | Alanine [-$Ni^0$] [+$NH_3$] (mM) | Alanine [+$Ni^0$] [-$NH_3$] (mM) | Alanine [+$Ni^0$] [+$NH_3$] (mM) |
|---|---|---|---|---|
| [1] | n.d. | n.d. | n.d. | 1.4762 |
| [2] | n.d. | n.d. | n.d. | 1.2491 |
| [3] | n.d. | n.d. | n.d. | 1.1224 |

| | Lactate [-$Ni^0$] [-$NH_3$] (mM) | Lactate [-$Ni^0$] [+$NH_3$] (mM) | Lactate [+$Ni^0$] [-$NH_3$] (mM) | Lactate [+$Ni^0$] [+$NH_3$] (mM) |
|---|---|---|---|---|
| [1] | n.d. | n.d. | 9.7135 | 9.3287 |
| [2] | n.d. | n.d. | 10.4968 | 8.3805 |
| [3] | n.d. | n.d. | 9.7113 | 7.6948 |

| | Acetate [-$Ni^0$] [-$NH_3$] (mM) | Acetate [-$Ni^0$] [+$NH_3$] (mM) | Acetate [+$Ni^0$] [-$NH_3$] (mM) | Acetate [+$Ni^0$] [+$NH_3$] (mM) |
|---|---|---|---|---|
| [1] | 0.0148 | 0.1617 | 0.8537 | 0.3320 |
| [2] | n.d. | 0.1906 | 0.8837 | 0.2974 |
| [3] | 0.0554 | 0.137 | 0.9038 | 0.3017 |

Supplemental Figure 2. **Raw data Figure 1.** Raw data of product concentrations with and without nickel and ammonium chloride. Initial concentrations were 20 mM pyruvate and 200 mM ammonium chloride. Ni-$SiO_2$/$Al_2O_3$ (1 mmol of metal atoms) was added as solid phase powder in a total reaction volume of 1.5 mL. The reaction was performed under a 5 bar Ar atmosphere, initial pH 11 with KOH, the reaction time was 18 h at 100°C. No $H_2$ was added. Reactions were performed in triplicates.

| pH | Pyruvate [1] (mM) | Pyruvate [2] (mM) | Pyruvate [3] (mM) | Lactate [1] (mM) | Lactate [2] (mM) | Lactate [3] (mM) |
|---|---|---|---|---|---|---|
| **7** | n.d. | n.d. | n.d. | 11.3001 | 10.6799 | 10.8950 |
| **8** | n.d. | n.d. | n.d. | 13.4924 | 16.1508 | 15.1491 |
| **9** | n.d. | n.d. | n.d. | 17.1124 | 17.4198 | 17.0749 |
| **10** | n.d. | n.d. | n.d. | 14.5391 | 16.2807 | 15.7643 |
| **11** | n.d. | n.d. | n.d. | 9.7478 | 10.9123 | 10.0172 |

| pH | Acetate [1] (mM) | Acetate [2] (mM) | Acetate [3] (mM) |
|---|---|---|---|
| **7** | 0.1191 | 0.0973 | 0.0879 |
| **8** | 0.1281 | 0.2965 | 03396 |
| **9** | 0.2964 | 0.4132 | 0.3113 |
| **10** | 0.1296 | 0.5342 | 0.7850 |
| **11** | 0.2322 | 0.2650 | 0.2455 |

Supplemental Figure 3. **Raw data Figure 2.** Raw data of lactate product concentrations. The pH was set to 7, 8, 9, 10, and 11 with KOH, respectively, and the reaction time was set to 2 h. Other reaction parameters are as in Figure 1.

Raw data Figure 3A [1h]

| Temperature (ºC) | Pyruvate [1] (mM) | Pyruvate [2] (mM) | Pyruvate [3] (mM) | Lactate [1] (mM) | Lactate [2] (mM) | Lactate [3] (mM) |
|---|---|---|---|---|---|---|
| **25** | 9.8090 | 11.0657 | 11.2670 | n.d. | n.d. | n.d. |
| **40** | 13.1949 | 15.2201 | 12.0138 | n.d. | n.d. | n.d. |
| **60** | 13.4307 | 15.4054 | 12.9687 | n.d. | n.d. | n.d. |
| **80** | 12.4268 | 14.1875 | 11.5273 | 1.0875 | 0.7280 | 1.0111 |
| **100** | 0.2891 | 0.0308 | 0.0592 | 14.6500 | 13.1054 | 14.0402 |

| Temperature (ºC) | Acetate [1] (mM) | Acetate [2] (mM) | Acetate [3] (mM) |
|---|---|---|---|
| **25** | 0.0164 | 0.0155 | 0.0156 |
| **40** | 0.0567 | 0.0580 | 0.0497 |
| **60** | 0.0917 | 0.1231 | 0.0926 |
| **80** | 0.1022 | 0.1334 | 0.0776 |
| **100** | 0.1115 | 0.0949 | 0.0835 |

Raw data Figure 3B [2h]

| Temperature (ºC) | Pyruvate [1] (mM) | Pyruvate [2] (mM) | Pyruvate [3] (mM) | Lactate [1] (mM) | Lactate [2] (mM) | Lactate [3] (mM) |
|---|---|---|---|---|---|---|
| **25** | 14.4576 | 13.7264 | 15.4781 | n.d. | n.d. | n.d. |
| **40** | 17.0321 | 14.7814 | 13.8389 | n.d. | n.d. | n.d. |
| **60** | 15.4497 | 14.9801 | 15.9499 | n.d. | n.d. | n.d. |
| **80** | 1.0500 | 0.6079 | 1.1167 | 12.7927 | 13.8431 | 13.9612 |
| **100** | n.d. | n.d. | n.d. | 17.1124 | 17.4198 | 17.0749 |

| Temperature (ºC) | Acetate [1] (mM) | Acetate [2] (mM) | Acetate [3] (mM) |
|---|---|---|---|
| **25** | 0.0155 | 0.0155 | 0.0230 |
| **40** | 0.0529 | 0.0738 | 0.0417 |
| **60** | 0.0600 | 0.0553 | 0.0731 |
| **80** | 0.1001 | 0.1059 | 0.1116 |
| **100** | 0.2964 | 0.4132 | 0.3113 |

Raw data Figure 3C [4h]

| Temperature (ºC) | Pyruvate [1] (mM) | Pyruvate [2] (mM) | Pyruvate [3] (mM) | Lactate [1] (mM) | Lactate [2] (mM) | Lactate [3] (mM) |
|---|---|---|---|---|---|---|
| **25** | 12.0511 | 11.3594 | 12.2813 | n.d. | n.d. | n.d. |
| **40** | 13.3314 | 12.6450 | 12.0363 | n.d. | n.d. | n.d. |
| **60** | 10.4015 | 10.8549 | 12.3827 | 1.7391 | 1.7932 | 1.8348 |
| **80** | n.d. | n.d. | n.d. | 14.4273 | 12.3039 | 14.5249 |
| **100** | n.d. | n.d. | n.d. | 15.8362 | 18.0540 | 15.6835 |

| Temperature (ºC) | Acetate [1] (mM) | Acetate [2] (mM) | Acetate [3] (mM) |
|---|---|---|---|
| **25** | 0.0226 | 0.0275 | 0.0341 |
| **40** | 0.1178 | 0.1007 | 0.0850 |
| **60** | 0.0994 | 0.0767 | 0.0947 |
| **80** | 0.0478 | 0.0562 | 0.0316 |
| **100** | 0.1769 | 0.2044 | 0.1047 |

Raw data Figure 3D [18h]

| Temperature (ºC) | Pyruvate [1] (mM) | Pyruvate [2] (mM) | Pyruvate [3] (mM) | Lactate [1] (mM) | Lactate [2] (mM) | Lactate [3] (mM) |
|---|---|---|---|---|---|---|
| **25** | 13.2098 | 13.7484 | 12.7345 | n.d. | n.d. | n.d. |
| **40** | 13.7183 | 14.1686 | 14.3055 | n.d. | n.d. | n.d. |
| **60** | 14.3702 | 11.0668 | 11.7109 | 0.3487 | 0.2389 | 0.2656 |
| **80** | n.d. | n.d. | n.d. | 12.7042 | 11.7315 | 11.4080 |
| **100** | n.d. | n.d. | n.d. | 11.0396 | 11.0131 | 12.1422 |

| Temperature (ºC) | Acetate [1] (mM) | Acetate [2] (mM) | Acetate [3] (mM) |
|---|---|---|---|
| **25** | 0.0216 | 0.0010 | 0.0149 |
| **40** | 0.3995 | 0.4563 | 0.3244 |
| **60** | 0.4048 | 0.3581 | 0.3950 |
| **80** | 0.2035 | 0.1508 | 0.1576 |
| **100** | 0.7373 | 0.8290 | 0.8542 |

Supplemental Figure 4. **Raw data Figure 3.** Raw data of lactate product concentrations. Parameters are as in Figure 1 unless otherwise indicated. The reaction time was 1h, 2h, 4h, and 18 h. Temperature was 25ºC, 40ºC, 60ºC, 80ºC, and 100ºC. The pH was set to 9 with KOH.

| $Ni-SiO_2/Al_2O_3$ (mM) | Pyruvate [1] (mM) | Pyruvate [2] (mM) | Pyruvate [3] (mM) | Lactate [1] (mM) | Lactate [2] (mM) | Lactate [3] (mM) |
|---|---|---|---|---|---|---|
| **0.05** | 13.1142 | 15.3396 | 12.1251 | 1.3007 | 1.1211 | 1.4426 |
| **0.11** | 12.5910 | 11.9418 | 10.4562 | 3.1723 | 3.3194 | 3.0858 |
| **0.22** | 11.2084 | 10.7782 | 11.4425 | 4.2738 | 4.2992 | 4.9393 |
| **0.33** | 7.5484 | 7.8566 | 9.5385 | 6.0045 | 7.4879 | 6.9266 |
| **0.66** | 1.4955 | 1.7374 | 2.4170 | 14.9149 | 14.2723 | 12.5128 |
| **1** | n.d. | n.d. | n.d. | 17.4198 | 17.0749 | 17.1124 |
| **1.5** | n.d. | n.d. | n.d. | 17.6590 | 16.0863 | 16.5755 |

| $Ni-SiO_2/Al_2O_3$ (mM) | Acetate [1] (mM) | Acetate [2] (mM) | Acetate [3] (mM) |
|---|---|---|---|
| **0.05** | 0.5562 | 0.8460 | 0.6070 |
| **0.11** | 0.1510 | 0.1342 | 0.0922 |
| **0.22** | 0.1483 | 0.1821 | 0.2157 |
| **0.33** | 0.1348 | 0.1464 | 0.1534 |
| **0.66** | 0.1731 | 0.1403 | 0.1361 |
| **1** | 0.4132 | 0.3113 | 0.2964 |
| **1.5** | 0.1464 | 0.1467 | 0.1197 |

Supplemental Figure 5. **Raw data Figure 4.** Raw data of lactate product concentrations. Parameters are as in Figure 1 unless otherwise indicated. $Ni-SiO_2/Al_2O_3$ was added as solid phase at a concentration of 0.05, 0.11, 0.22, 0.33, 0.66, 1, and 1.5 mmol, respectively. The pH was 9 (KOH), and reaction time was 2 h. Each reaction was performed in triplicates.

| | 4-Methyl-2-oxopentanoate (mM) | 2-Hydroxyisocaproate (mM) |
|---|---|---|
| [1] | 0.2931 | 19.5348 |
| [2] | 0.7288 | 19.5591 |
| [3] | 0.7584 | 19.3958 |

| | 3-Methyl-2-oxopentanoate (mM) | 2-Hydroxy-3-methylvalerate (mM) |
|---|---|---|
| [1] | 0.9678 | 12.4926 |
| [2] | 0.8970 | 13.4035 |
| [3] | 0.6670 | 12.6702 |

| | 2-Oxoglutarate (mM) | 2-Hydroxyglutarate (mM) |
|---|---|---|
| [1] | n.d. | 21.7220 |
| [2] | n.d. | 20.1870 |
| [3] | n.d. | 20.3880 |

| | Fumarate (mM) | Succinate (mM) |
|---|---|---|
| [1] | n.d. | 24.2600 |
| [2] | n.d. | 24.3900 |
| [3] | n.d. | 24.7000 |

Supplemental Figure 6. **Raw data Figure 5.** Raw data of reduction product concentrations. Parameters are as in Figure 1 unless otherwise indicated. Educt concentration was set to 20 mM. The reaction time was set to 2 h, and pH was set to 9 with KOH

| | 4-Methyl-2-oxopentanoate (mM) | Leucine (mM) | 2-Hydroxyisocaproate (mM) |
|---|---|---|---|
| [1] | n.d. | 1.3336 | 11.6780 |
| [2] | n.d. | 2.2125 | 11.8850 |
| [3] | n.d. | 1.3001 | 12.1904 |

| | 3-Methyl-2-oxopentanoate (mM) | Isoleucine (mM) | 2-Hydroxy-3-methylvalerate (mM) |
|---|---|---|---|
| [1] | n.d. | 0.9117 | 8.0438 |
| [2] | n.d. | 1.2431 | 7.5011 |
| [3] | n.d. | 0.7421 | 6.4658 |

| | 2-Oxoglutarate (mM) | 2-Hydroxyglutarate (mM) | Glutamate (mM) | 2-Oxoproline (mM) |
|---|---|---|---|---|
| [1] | n.d. | 1.7629 | n.d. | 11.8812 |
| [2] | n.d. | 2.6339 | n.d. | 11.4198 |
| [3] | n.d. | 1.4145 | n.d. | 11.1556 |

Supplemental Figure 7. **Raw data Figure 6.** Raw data of reductive amination product concentrations. Parameters are as in Figure 1 unless otherwise indicated. The reaction time was 72 h.
